# Assessing the effects of acute temperature changes of sea water temperature due to climate change on Scottish farmed Atlantic salmon and investigating genetic mitigation

**DOI:** 10.1101/2024.01.29.577726

**Authors:** Smaragda Tsairidou, Jarl van den Berg, Stephen Tapping, Halina Sobolewska, Alastair Hamilton

## Abstract

Changes in growth, survival, maturation and health in Atlantic salmon post-smolts were observed under moderate and more extreme heat-wave conditions for the west coast of Scotland. A Scottish Atlantic salmon breeding programme population of 518 salmon of age 13-14 months, was observed for 4 weeks in summer, in tanks with ambient water temperature or ∼ 4 degrees and ∼ 8 degrees above ambient temperature. Data were recorded for the fish before and after the challenge. All fish were genotyped using a custom SNP (single nucleotide polymorphism) array (8,978 SNP genotypes after quality control). Temperature-dependant genotype-by-environment interactions and the potential for selective breeding to improve resilience were investigated. Statistical analyses revealed significant differences between thermal environments for body weight, average daily weight gain, and survival, while higher temperature induced earlier maturation, and an increase of gill health scores. QPCR (quantitative polymerase chain reaction) analyses revealed the presence of *Aeromonas hydrophila*, a pathogen typically more prevalent in warmer climates. Using genomic relationships and animal mixed models, body weight and average daily weight gain provided moderate heritabilities, while between-tank genetic correlations were close to 1, indicating no significant re-ranking of genotypes between the different thermal environments. These findings suggest that even short-term exposure to heat stress may be sufficient to negatively impact survival and gill health, and induce earlier maturation. However, observations took place within a commercial farm where having replicates for each thermal environment was not possible, hence further experiments with larger populations, exposed to more prolonged heat stress are needed.

## 1. Introduction

Extreme weather events such as heat waves are increasing in duration, frequency and intensity due to climate change (Russo et al. 2014; IPCC 2023; Klingelhöfer et al. 2023). Strong evidence supports that those changes are linked to anthropogenic influences on global climate (Coumou and Rahmstorf 2012), such as greenhouse gas emissions, which have resulted in climatic warming at unprecedented rates (IPCC 2021). Climate change is an emerging challenge for the aquaculture sector as seafood production is vulnerable to interactions with environmental factors. Fish provide 3.3 billion people with over 20% of their average per capita animal protein intake, and is a major contributor to global food security, especially for low and middle income countries (FAO 2020). In the UK, salmonid aquaculture is a major contributor to the national and rural economies, with Atlantic salmon farmed in Scotland being the UK’s largest fresh food export worth £614 million (Scottish Government 2022), and the UK being the third largest producer in the world, following Norway and Chile.

Salmonids are ectotherms and each species can survive and optimally perform within a range of temperatures. Atlantic salmon has a growth range between 6°C and 22.5°C, with maximum growth at 16°C (Lightfoot 2008). Therefore, heat waves can have an impact on salmon performance, health and welfare, can affect the cost of production, and can generate shifts in the geographic regions where salmon farming is feasible and profitable. For areas such as Norway, Tasmania, Atlantic Canada, and Scotland, where salmon populations are relatively naïve to higher seawater temperatures, heat-stress conditions may have detrimental effects on their health, welfare and performance, and can result in infectious disease outbreaks of increased severity and frequency. Therefore, understanding the possible impacts of climate change on salmon production is crucial for improving resilience and designing effective mitigation strategies to ensure profitability and long-term sustainability of the sector.

Previous studies in salmonids suggest the existence of phenotypic and possibly also genetic variation in their thermal tolerance and response to heat stress, affecting several commercially important production and health traits. Between-family differences in upper temperature tolerance in Atlantic salmon have been measured via the critical maximum temperature (Anttila et al. 2013), and the incremental thermal maximum (Ignatz et al. 2023). Debes et al. (2021) reported an antagonistic relationship between growth and thermal tolerance, highlighting the potential role of genotype-by-environment interactions and specifically the role of increasing acclimation temperature in reducing between-population variation for thermal tolerance. In addition, increased mortality rates associated with higher water temperature have been reported in Sockeye salmon populations (Eliason et al. 2011), and in rainbow trout (Chen et al. 2015). Stress conditions can compromise the effectiveness of the immune response, and heat-stress has been associated with a wide range of pathologies, in particular related to gill health, including earlier and more severe pathology in *Paramoeba perurans* infection in Atlantic salmon (Benedicenti et al. 2019). Further, early onset of sexual maturation in Atlantic salmon has been found to be controlled by both genetic and environmental factors (Wild et al. 1994). Reversible phenotypic plasticity via acclimation is known to allow adaptation to some extent, however, previous studies in salmonids have shown a limited capacity to substantially increase their thermal tolerance via this mechanism (Elliott 1991; Baroudy and Elliott 1994). In contrast, after several generations of exposure and natural selection, adaptation of metabolic and physiological mechanisms to better cope with higher temperatures has been observed in a rainbow trout strain (Chen et al. 2015), while in blue tilapia, differential gene responses have been associated with cold water tolerance (Nitzan et al. 2019).

Selective breeding has been increasingly implemented by the aquaculture industry, and breeding programmes have been developed for Atlantic salmon and other aquaculture species to improve economically important production and health traits. For example, research in salmon genetics has allowed growth improvements (Gjedrem and Rye 2018), and identification of genomic variants that impact resistance to important infectious diseases (e.g. Houston et al. 2010). By using advanced computational techniques such as genotype imputation, large-scale genomic data required for selective breeding can be obtained in a cost-effective manner, rendering the process accessible for more breeders and producers (Tsairidou et al. 2020). However, to routinely include robustness to climate change effects into the breeding goals, we would need to identify suitable phenotypes (i.e., target traits), and detect genetic variation associated with those traits.

Using genomic data to estimate genetic correlations between different, e.g. thermal, environments allows detection of genotype re-ranking due to genotype-by-environment interactions (Sae-Lim et al. 2015). This is pertinent, firstly because it shows that different genotypes vary in their relative performance in different environments, and secondly because, if unaccounted for, such interactions could cause selective breeding in current environments to be less effective in future environments affected by climate change. The high fecundity of salmonids allows for large full sibling families to be available in typical aquaculture breeding programmes, hence it allows testing of full siblings in different environments, enabling powerful assessments of genotype-by-environment interactions and phenotypic plasticity.

With a focus on short-term temperature spikes, the aims of this study were to (i) record and phenotypically assess changes in growth, survival, maturation, and health traits in salmon due to moderate and more extreme heat-wave conditions, in order to identify commercially important traits that may be affected by heat waves in Scottish waters, and (ii) estimate genetic variation and genetic correlations between different thermal environments, to investigate temperature-dependant genotype-by-environment interactions. Specifically, the effects of heat stress on post-smolts were observed within a commercial farm, in two different thermal environments which resembled: (a) a moderate heat wave at the west coast of Scotland, which is expected to become more frequent due to climate change, leading to water temperatures just above salmon’s optimum growth temperature, and (b) a more extreme heat-wave event, generating water temperatures at the upper edge of salmon’s growth range.

## 2. Materials and methods

### 2.1 Phenotypes

A commercial Scottish Atlantic salmon breeding programme population, comprising 518 seawater adapted salmon post-smolts, 242 female and 276 male, of age 13-14 months, were exposed to three different thermal environments for 4 weeks during July 2021. The tanks were 2×3m^3^ with partition, using recirculating water. In Tank A, water was at ambient temperature, in Tank B, the water was heated to 4 degrees above the ambient temperature, and in Tank C, the water was heated to 8 degrees above ambient (Figure 1). Tank B represented heat wave conditions on the west coast of Scotland, which are expected to occur more frequently due to climate change, and, Tank C represented a more dramatic temperature increase as expected in future heat wave events. The fish were acclimatised for 3 months. During the challenge, all fish in all tanks received the same amount and type of feed.

**Figure 1.**
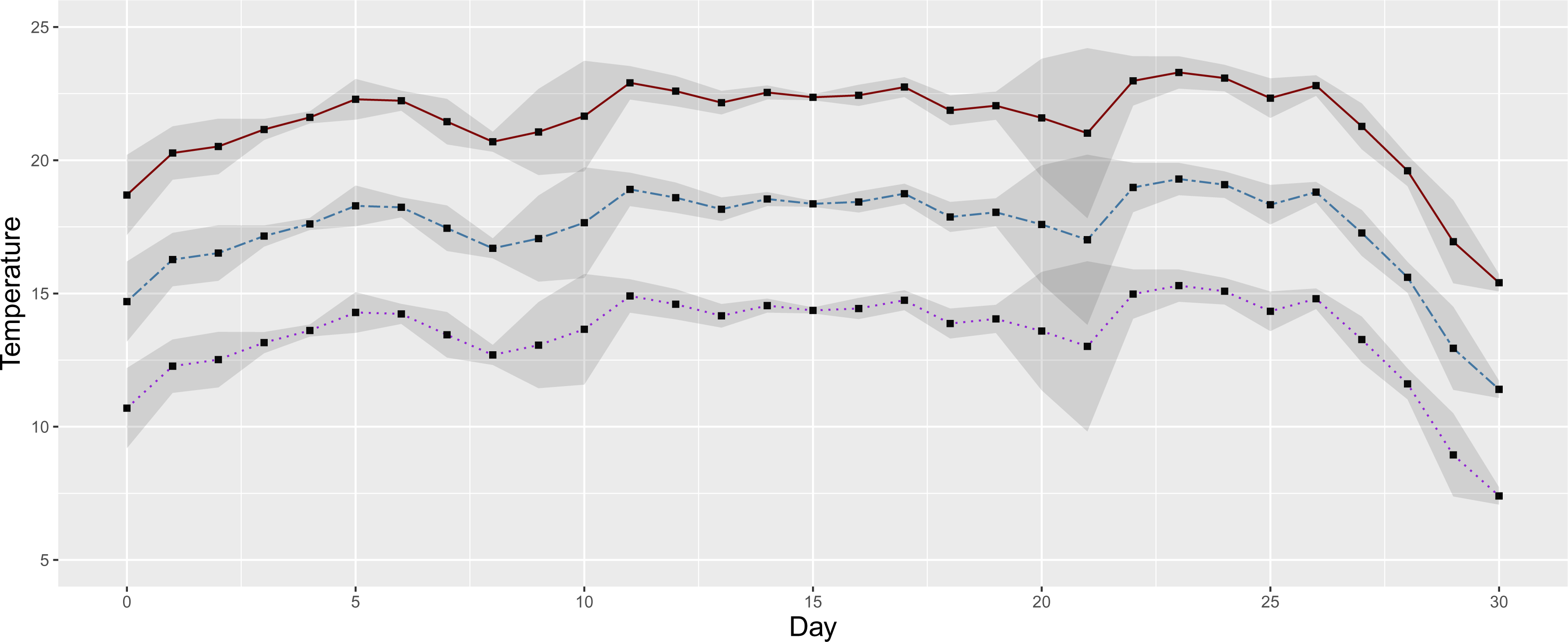
Mean daily temperature in each of the three tanks i.e. (i) Tank A (purple dotted line): the water was at ambient temperature, (ii) Tank B (blue dashed line): the water was heated to 4 degrees above the ambient temperature, and, (iii) Tank C (red solid line): the water was heated to 8 degrees above ambient. The grey shaded ribbon represents the standard deviation

The water temperature and the amount of dissolved Oxygen (in mg/L and % saturation) were recorded for the high temperature tank every 10 minutes, using the OxyGaurd system. Mean daily temperature started at 18.7°C at the beginning of the challenge, then gradually increased, and reached a maximum of 23.3°C on day 23 (Figure 1); for the greatest part of the observation period the mean daily temperature in Tank C was > 20°C. The minimum mean oxygen concentration was 7.6 mg/l and the maximum was 10.1 mg/l; the minimum and maximum % oxygen saturation were 104 and 131.8 respectively (Figure 2). Water salinity varied with the tide between 25 and 33ppt.

**Figure 2.**
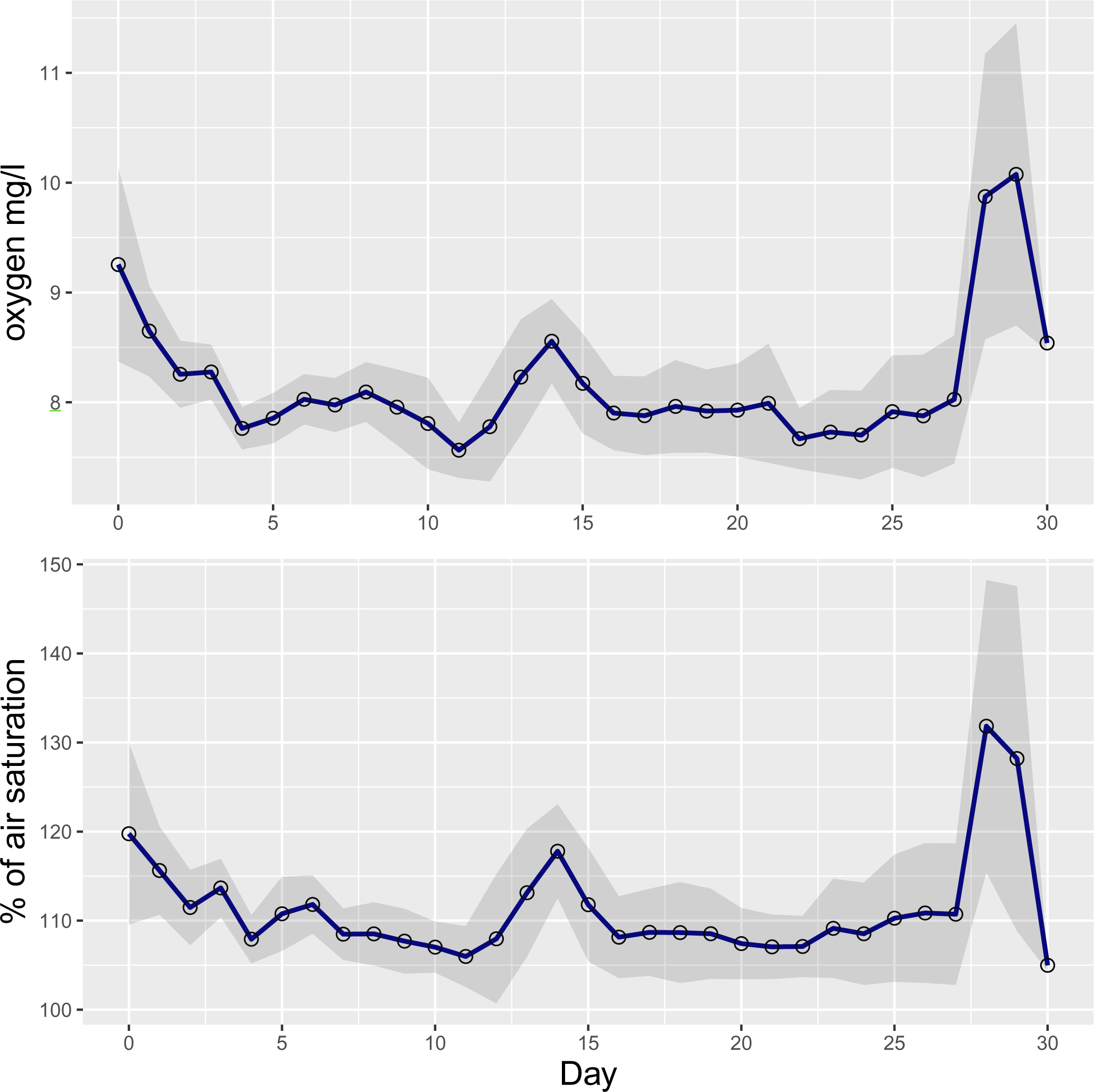
Mean daily oxygen concentration (mg/l), and mean daily oxygen % saturation

**Figure 3.**
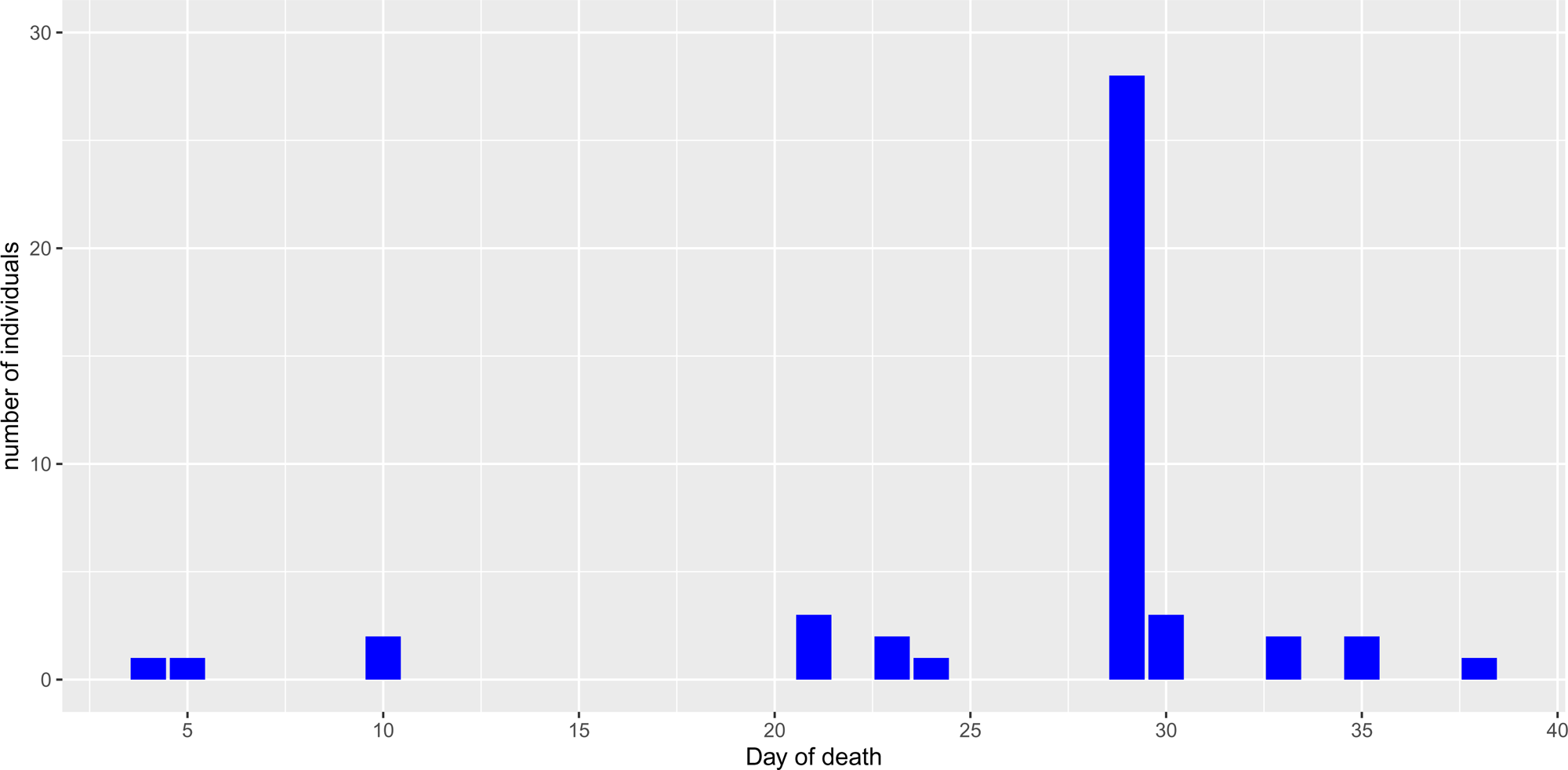
Distribution of mortalities during and soon after the challenge, where day of death was calculated from the recorded date of mortality as the number of days since 1/7/21; day 29 is the day of data capture after the challenge. The Y-axis shows the number of individuals that died on that day

Phenotypes were recorded as part of the standard operating procedure for routine data capture on the farm. The fish were sedated using tricaine, and body weight and length were measured before and after the challenge. After the challenge, assessment of maturation (grilse), ultrasound scanning, and gill health scoring were performed. In addition, gill swab samples were collected for QPCR analyses.

### 2.2 Statistical analysis of phenotypes

Statistical significance of differences between tanks were tested via one-way analysis of variance (ANOVA), and P-values were obtained using the F statistic and compared to the 0.05 significance level. Further, for body weight, an ANCOVA analysis was performed, adjusting for the before challenge weight fitted as a covariate in the analysis of after challenge weight, following the model: 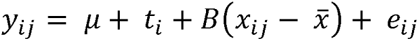, where *y_ij_* is body weight after the challenge for individual *j* in tank *i*, µ is the grand mean, *t_i_* is the tank, *x_ij_* is the body weight before challenge for individual *j* in tank *i* fitted as covariate, 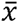 is the global mean for covariance *x*, *B* is the slope, and *e_ij_*the error term.

Average daily weight gain was calculated as the difference between body weight before and after the challenge, divided by the number of days between the two data captures (t = 28). To test the hypothesis that changes in temperature may affect the fish fusiform shape and hence the efficiency of their basic metabolic rate, the body-weight to body-length ratio (Fulton’s performance index) was calculated as *k* = 100 × (*w/l*^3^), where *w* is the body weight in grams and *l* is the body length in cm.

Maturation was recorded as binary (grilse/no grilse) at the after-challenge data capture (Table 1). Comparison between thermal environments was performed via Pearson’s chi-squared test. Mortalities were recorded daily during the challenge. In the first 10 days after the challenge, there was a small number of mortalities; those were included in subsequent analysis of survival data, as is likely that heat stress made the fish less tolerant to handling processes. Overall, 3 mortalities were recorded as physical damage, 28 as handling damage, and 15 had no obvious cause. Deformities were observed on the tail of only two fish; hence this phenotype was not pursued further in the analyses.

**Table 1.** Number of individuals with phenotypes, prevalence of grilse, mortalities, number of families with offspring in each tank, and number of individuals with genotypes.

| Tank | n<sub>phenotyped</sub> | Grilse | Mortalities | Families | n<sub>genotyped</sub> |
| --- | --- | --- | --- | --- | --- |
| A | 193 | 9 | 17 | 54 | 182 |
| B | 171 | 26 | 5 | 53 | 165 |
| C | 154 | 22 | 24 | 51 | 145 |
| Total | 518 | 57 | 46 | NA | 492 |

Gill health was assessed via observation of gross gill pathologies for fish in Tanks B and C, and scored using a multifactorial score (Fridman et al. 2021). Briefly, following Fridman et al. (2021) macroscopic pathological changes in the gills can be categorised as 0 (healthy), 1 (very light), 2 (more extensive), 3 (extensive), 4 (advanced), or 5 (severe). A screening QPCR assay was performed using 10 gill swab samples selected randomly from each of the Tanks B and C (n_total_ = 20). The selection of individual swabs was informed by the gill health scores observed in that tank so that all observed scores were represented in the QPCR (Table 2). Taqman MGB QPCR analysis was performed targeting the following gill pathogens: *Candidatus* Branchiomonas cysticola (Mitchell et al. 2013), *Candidatus* Piscichlamydia salmonis (Steinum et al. 2010), salmon gill pox virus (SGPV) (Gjessing et al. 2015), *Desmozoon lepeophtherii* (Nylund et al. 2010), *Neoparamoeba perurans* (Fringuelli et al. 2012), and *Neoparamoeba* spp (Fringuelli et al. 2012). Ct values were normalised against a Taqman assay targeting salmon elongation factor-α (Fringuelli et al. 2012). In addition, Eva Green QPCR assays were performed targeting *Flavobacterium branchiophilum*, *Aeromonas hydrophila* and total bacterial load (Steinum et al. 2010).

**Table 2.** Gill health scores and corresponding number of samples from each of the tanks.

| Tank | Gill health score | Number of samples |
| --- | --- | --- |
| B | 0 | 4 |
| B | 1 | 3 |
| B | 2 | 3 |
| C | 0 | 5 |
| C | 1 | 5 |

### 2.3 Genotypes and Quality Control

Genotypes were available for 8,981 SNPs obtained from fin-clip samples from 493 fish, genotyped using a custom SNP array. Quality Control was performed in Plink/1.90-beta4.1 (Purcell et al. 2007); individuals with a percentage of missing genotypes > 15 %, and SNPs with a minor allele frequency (MAF) < 5%, missing genotypes > 95%, or Hardy-Weinberg Equilibrium P-value < 10^-5^ were excluded. After quality control, 8,978 SNPs for 492 salmon were retained, and were used to calculate the genomic relationship matrices. In addition, pedigree information was available for all fish.

### 2.4 Genomic relationships and structure analysis

The genomic relationship matrix (GRM) was calculated from the GenABEL/R (“ibs” function) kinship matrix (Amin et al. 2007) using SNP marker data, and then multiplied by two and inverted (“solve”/R function) for use in ASReml (see below). The genomic relationship between animals *i* and *j* was given by:

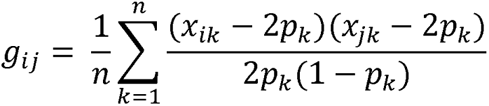

Where *g_ij_*are the genomic relationship between animals *i* and *j*, *n* is the number of loci used for estimating relationships; *x_ik_* is the count of alternative alleles (0, 1 or 2) of individual *i* at SNP locus k (the reference allele is arbitrarily chosen); and, *p_k_* is the frequency of the reference allele in the data. The diagonal elements are given by g_ii_ = 1+F_i_, where F_i_ is the inbreeding coefficient for individual *i*, calculated using SNP genotypes as follows:

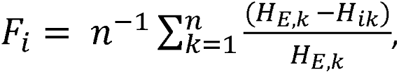

Where H_E,k_ is the expected heterozygosity at locus k assuming Hardy-Weinberg equilibrium, and H_ik_ is the observed heterozygosity for animal *i*.

Structure analysis using classical multi-dimensional scaling (“cmdscale” function in package “stats”, R/3.1.2) identified three distinct clusters in the data. After further exploration, the clusters were not found to be due to different sex of the fish, tank, or family (Supplementary Information 1), therefore principal components were fitted as covariates in all genetic models.

### 2.5 Heritability and genetic correlations

Firstly, heritability estimates were obtained across the entire population (n = 492) to obtain an indication of the genetic variation present in this dataset for the traits studied, namely: (a) directly recorded phenotypes in the three thermal environments, i.e. body weight and length before and after challenge, survival (dead/alive), maturation (grilse/non-grilse), and gill health score; and, (b) derived traits, i.e. the average daily weight gain calculated as the difference between body weight before and after the challenge, divided by the number of days between the two data captures (28 days), and, the day of death calculated from the recorded date of mortality, as the number of days since 1/7/21 (the date of data capture before the challenge). The following linear animal mixed model was fitted in ASReml/4.2, using either the genomic relationship matrix or pedigree relationships:

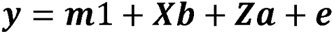

Where, **y** is the response variable (i.e. each of the traits described above); **m** is the overall mean; **b** is the vector of fixed effects associated with the incidence matrix **X**, comprising sex, tank, and the first two principal components estimated from the structure analysis fitted as covariates; **a** is the vector of random animal effects with **a**∼MVN(0, 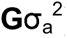) associated with the incidence matrix **Z**, where 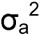 is the additive genetic variance; and, **e** is the residual error with **e**∼MVN(0, 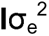). In addition, for survival and grilse, a generalized linear mixed animal model was fitted to take into account the binary character of these data, using the logit link function in ASReml which assumes an underlying logistic liability distribution.

Secondly, heritability estimates were obtained within-environment, using only data collected from each tank. Trait definitions and models were the same as described above, with the difference that tank was not fitted in the fixed effects. A series of bivariate analyses were performed in ASReml/v4.2 to identify suitable starting values and optimize the estimation of covariance components between the traits measured in the three environments, using the GRM calculated from all individuals. The response variables were re-defined such that, for example for body weight, three new traits were created, one for each thermal environment, where each individual had its body weight value if it was present in that tank and missing value otherwise. The principal components from the structure analyses were standardized by subtracting the mean and dividing by the standard deviation, and were fitted as covariates.

## 3. Results

### 3.1 Analyses of phenotypes

Phenotype data including body weight and length, average daily weight gain, Fulton’s performance index, grilse, survival, and gill health score were recorded for 518 salmon, in three different thermal environments (Table 1).

### 3.1.1 Body weight and length, average daily weight gain and Fulton’s performance index

Individual body weight and length ranged from 88 g to 476 g and from 200 mm to 380 mm respectively before challenge, and from 74 g to 618 g and from 210 mm to 510 mm after challenge. Mean and standard deviation for each tank is shown in Table 3.

**Table 3.** Phenotypic means (with standard deviation) before and after challenge for each tank (includes only survivors with weight record after the challenge).

| Trait | Tank A | Tank B | Tank C |
| --- | --- | --- | --- |
| Mean body weight (g) before challenge | 206 (70.2) | 195 (59.0) | 202 (67.7) |
| Mean body weight (g) after challenge | 280 (97.0) | 329 (81.6) | 315 (82.3) |
| Mean body length (mm) before challenge | 275 (28.8) | 270 (25.7) | 275 (28.2) |
| Mean body length (mm) after challenge | 301 (35.7) | 293 (27.6) | 294 (30.4) |
| Fulton's performance index before challenge | 0.96 (0.1) | 0.97 (0.2) | 0.95 (0.1) |
| Fulton's performance index after challenge | 1.00 (0.2) | 1.29 (0.1) | 1.23 (0.1) |
| Mean average daily weight gain (g per day) | 2.64 (1.4) | 4.78 (1.3) | 3.97 (1.2) |

There was a statistically significant difference between thermal environments for body weight (P-value < 0.05), which remained significant after adjusting for body weight before challenge. A moderate increase of water temperature, i.e. in Tank B with water temperature ∼ 4° higher than ambient, was beneficial for body weight compared to that in ambient temperature, but further exceeding salmon’s optimal zone with more extreme heat-stress in Tank C, had a negative impact on growth compared to Tank B, although still superior to Tank A (Figure 4).

**Figure 4.**
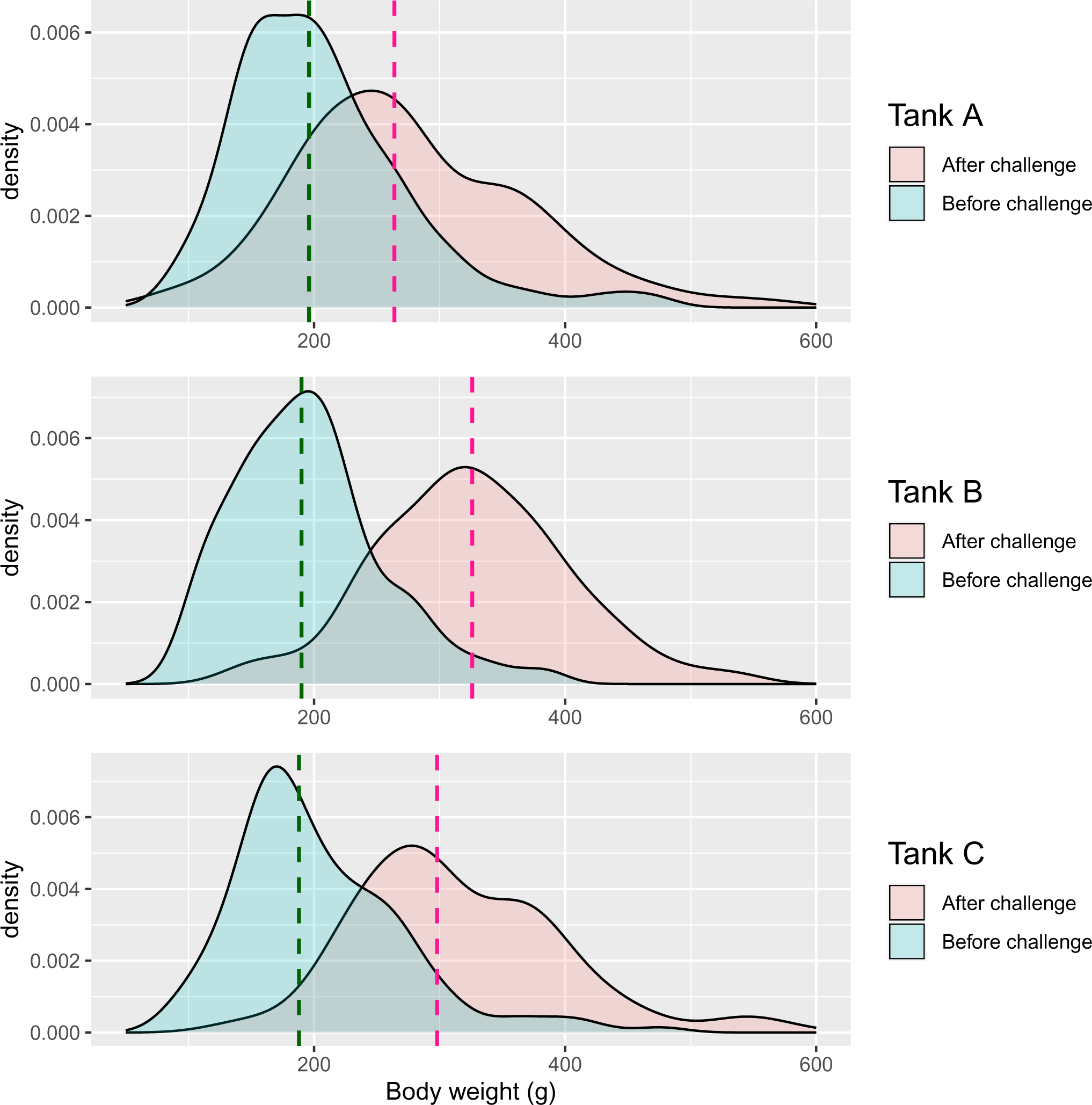
Distributions of body weight (g) before and after challenge for each environment; dashed lines represent the median before (green) and after (pink) challenge

This was consistent with observations for average daily weight gain, where higher temperature increased the average daily weight gain (Figure 5). Fish in Tank B performed better than fish in Tank C and this difference was statistically significant (pairwise comparison between tanks using the Tukey multiple comparisons of means test, 95% confidence level, P-value after adjustment for the multiple comparisons < 0.001). Normality assessment of the residuals for post-challenge weight from the ANCOVA showed only a minor departure from normality; nevertheless, the ANCOVA was repeated with the logarithmically transformed weights, and the difference between tanks remained significant. No significant differences for body length were found between thermal environments (Figure 6).

**Figure 5.**
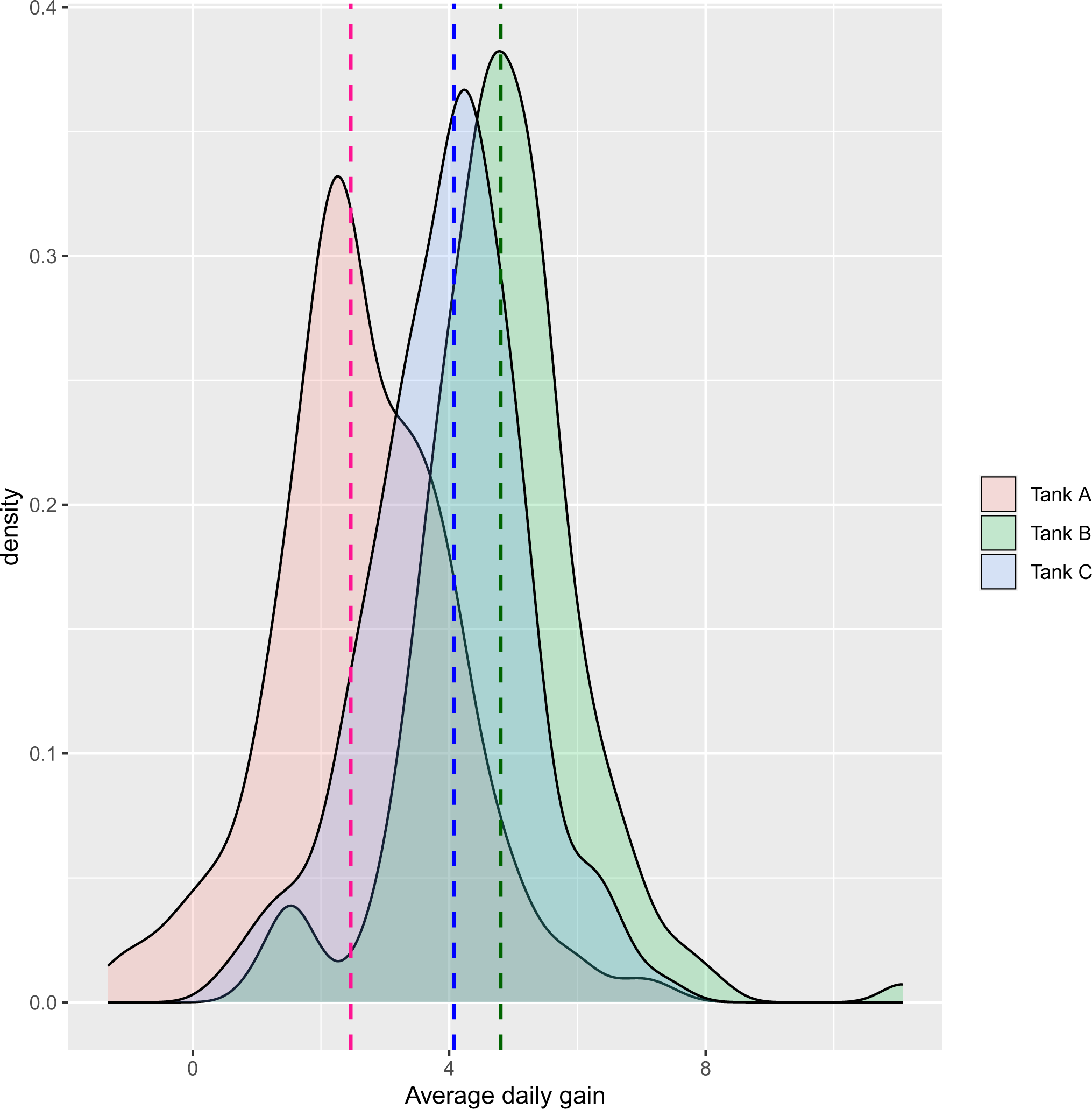
Distributions of average daily weight gain (g per day) in each of the three tanks; dashed lines represent the median for Tank A (blue), Tank B (pink) and Tank C (green)

**Figure 6.**
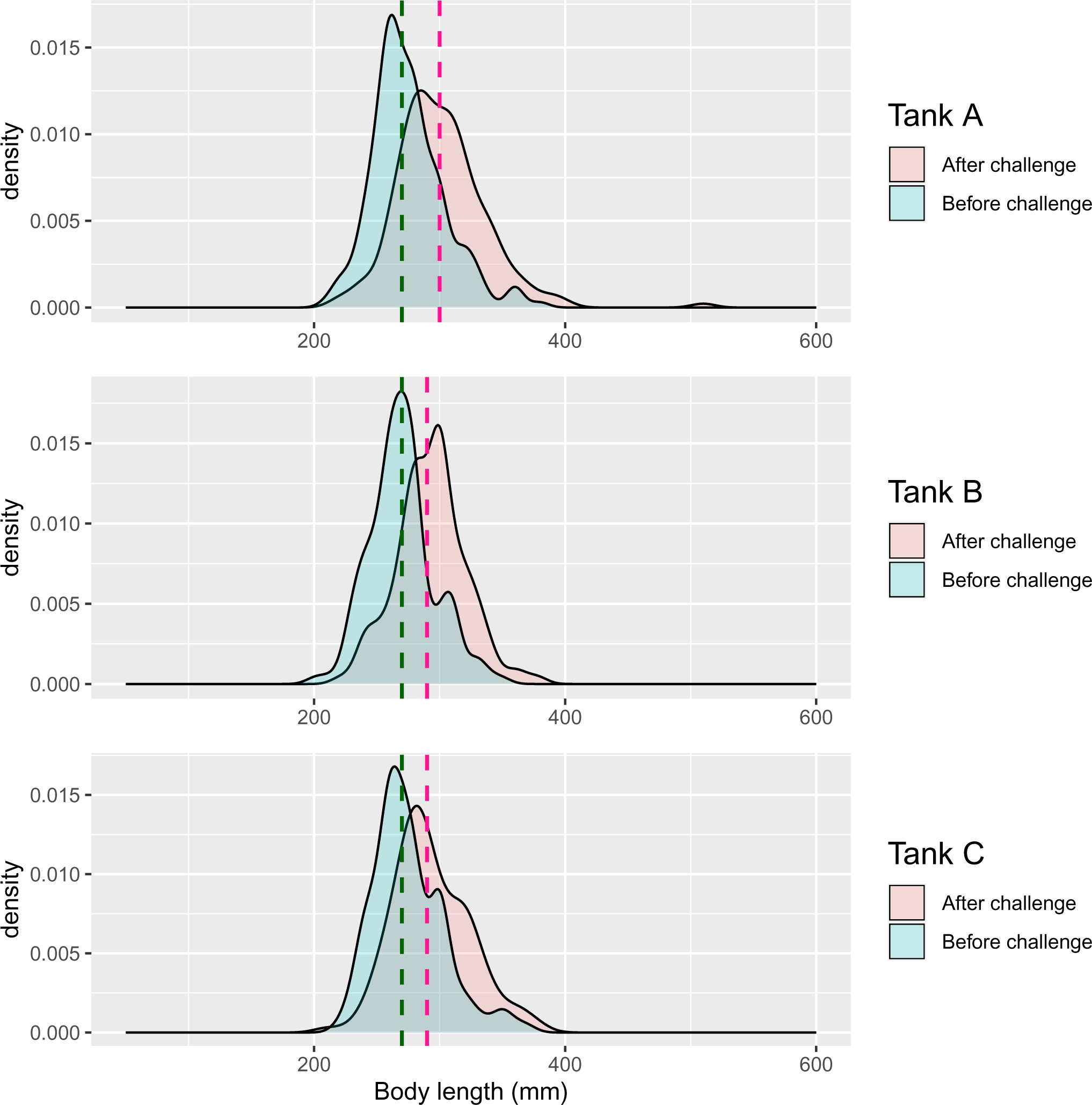
Distributions of body length (mm) before and after challenge for each environment; dashed lines represent the median before (green) and after (pink) challenge

In agreement with what was observed for body weight and average daily weight gain, the Fulton’s performance index was highest in Tank B, and then declined again for higher temperatures in Tank C (Figure 7), with Tanks B and C being higher than Tank A.

**Figure 7.**
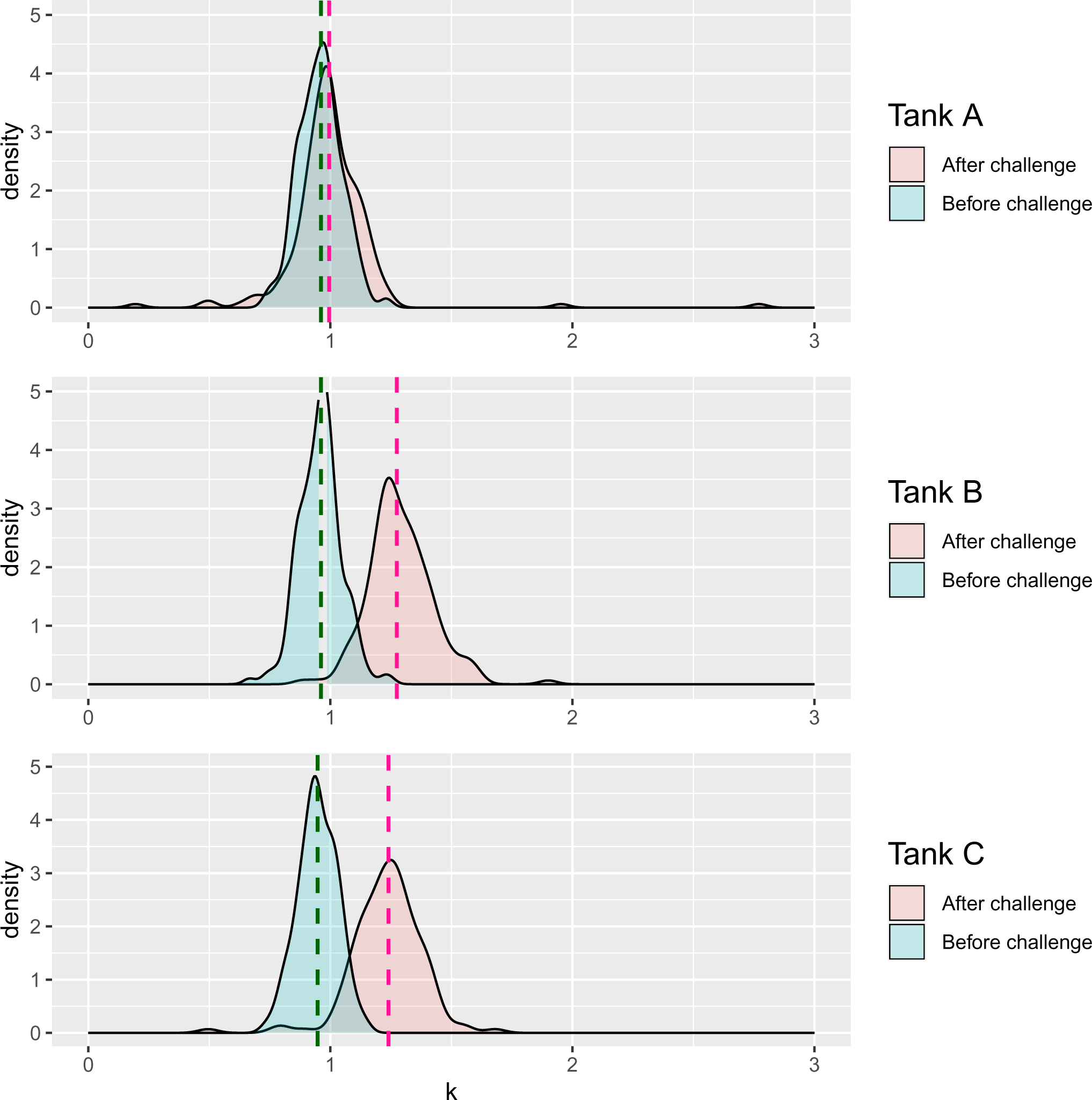
Fulton’s performance index k. The dashed lines represent the median before (green) and after (pink) challenge

#### 3.1.2 Maturation and survival

Maturation was assessed based on body colour, palpation of gonads and ultrasound scanning (Table 1). Comparison of maturation between thermal environments was performed via Pearson’s chi-squared test and Fisher’s exact test, and showed statistically significant effects for Tanks B and C (X-square test P-value < 0.01; Fisher’s exact test P-value < 0.001). Fish in higher temperatures had a higher rate of earlier maturation (Figure 8).

**Figure 8.**
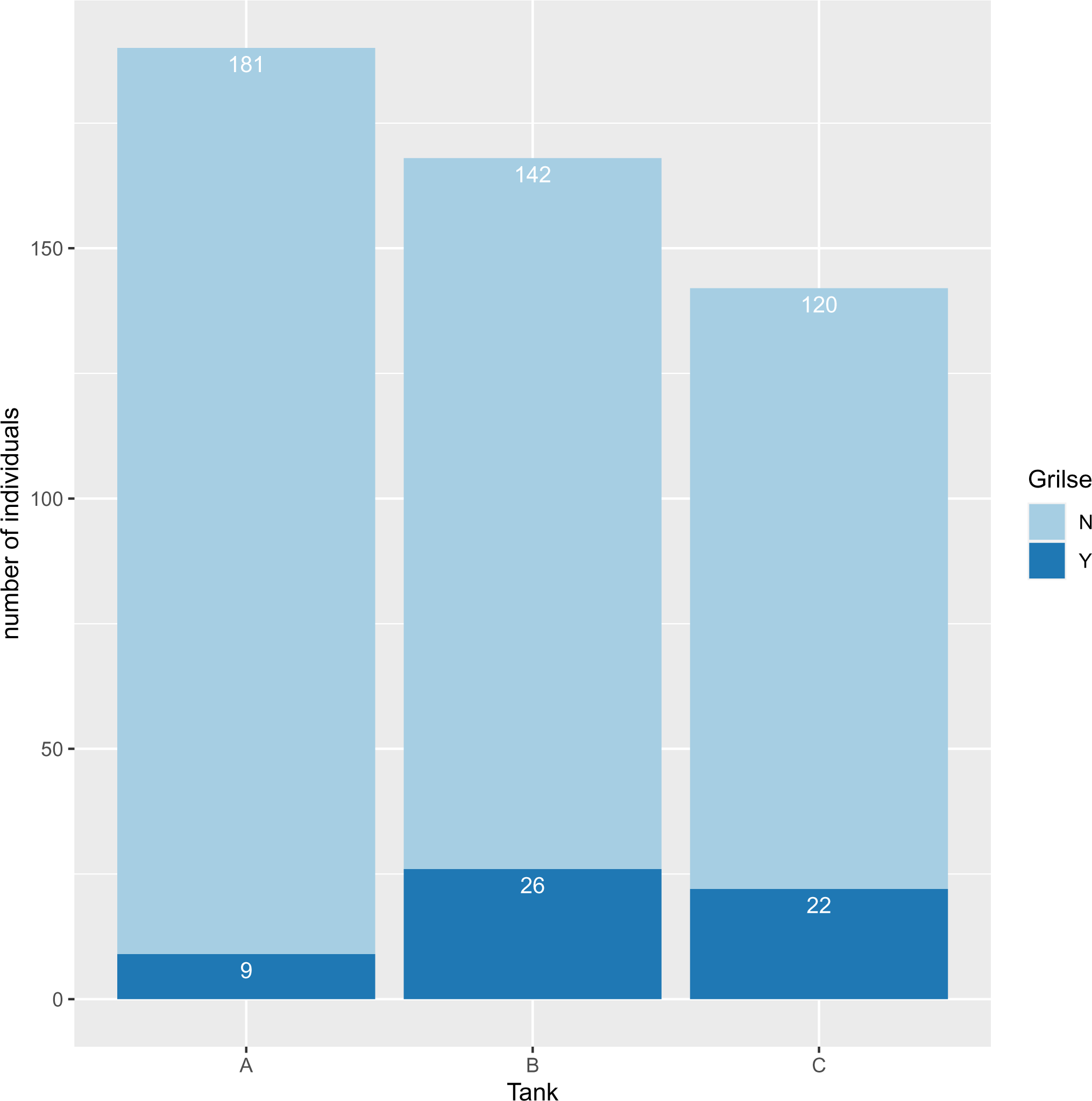
Bar plot of maturation indicator (grilse) in the different thermal environments, where n the number of individuals

10 mortalities occurred during the challenge, the majority of mortalities occurred on the day of data capture after the challenge, while 8 mortalities occurred during the first 10 days after the challenge (Table 1; Figure 3). There was a significant relationship between thermal environment and survival as indicated via the Pearson’s chi-squared test (P-value < 0.001) (Figure 9).

**Figure 9.**
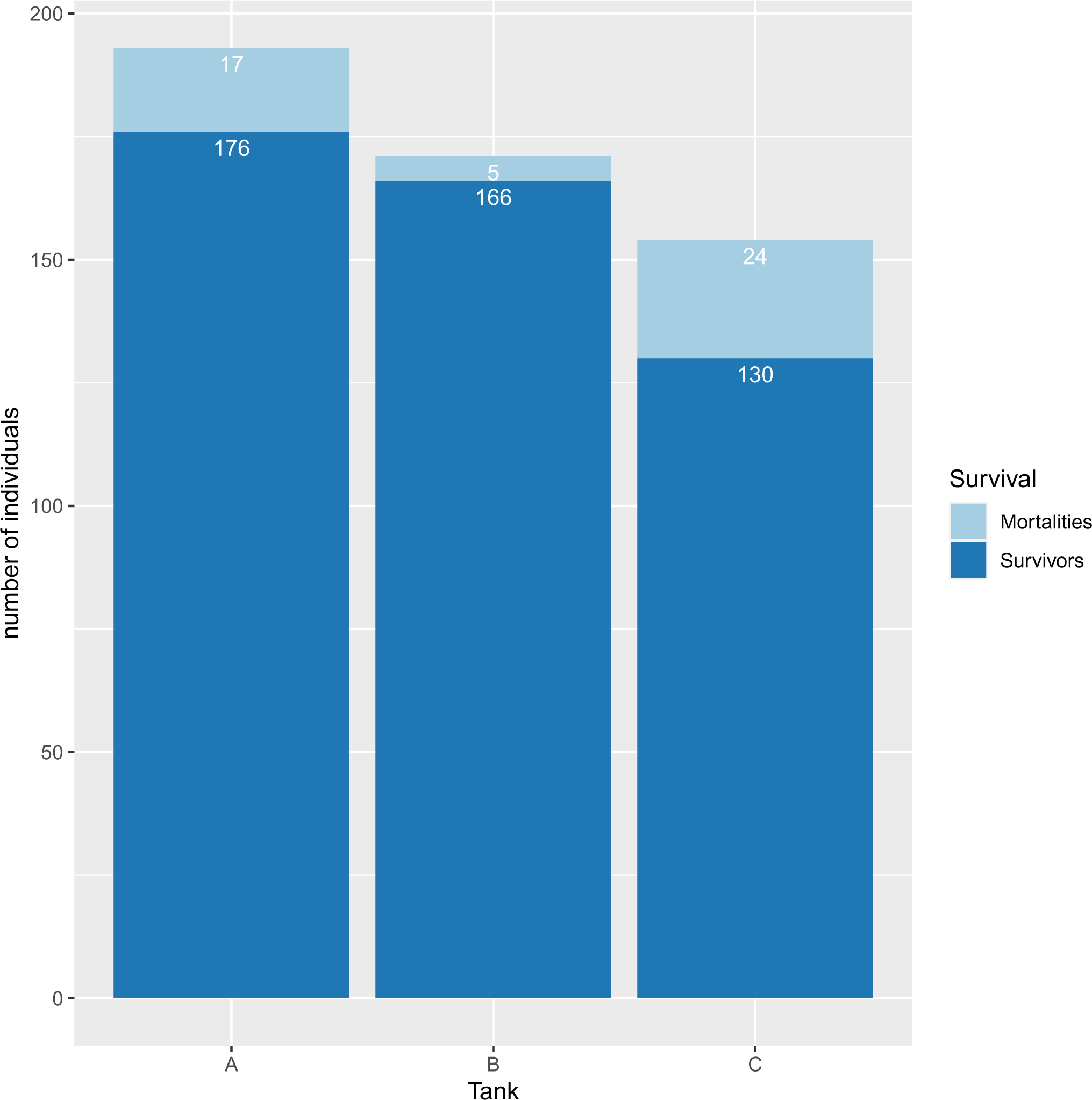
Comparison of mortalities between tanks, where n the number of individuals

#### 3.1.3 Gill health

Gill health scores were higher for fish exposed to higher temperatures, with scores 1 and 2 being notably more prevalent in tanks B and C (Figure 10) compared to the weekly scores routinely recorded during the same time-period and which indicated complete absence of gill lesions in this farm. The difference between tanks was statistically significant (P-value < 0.001).

**Figure 10.**
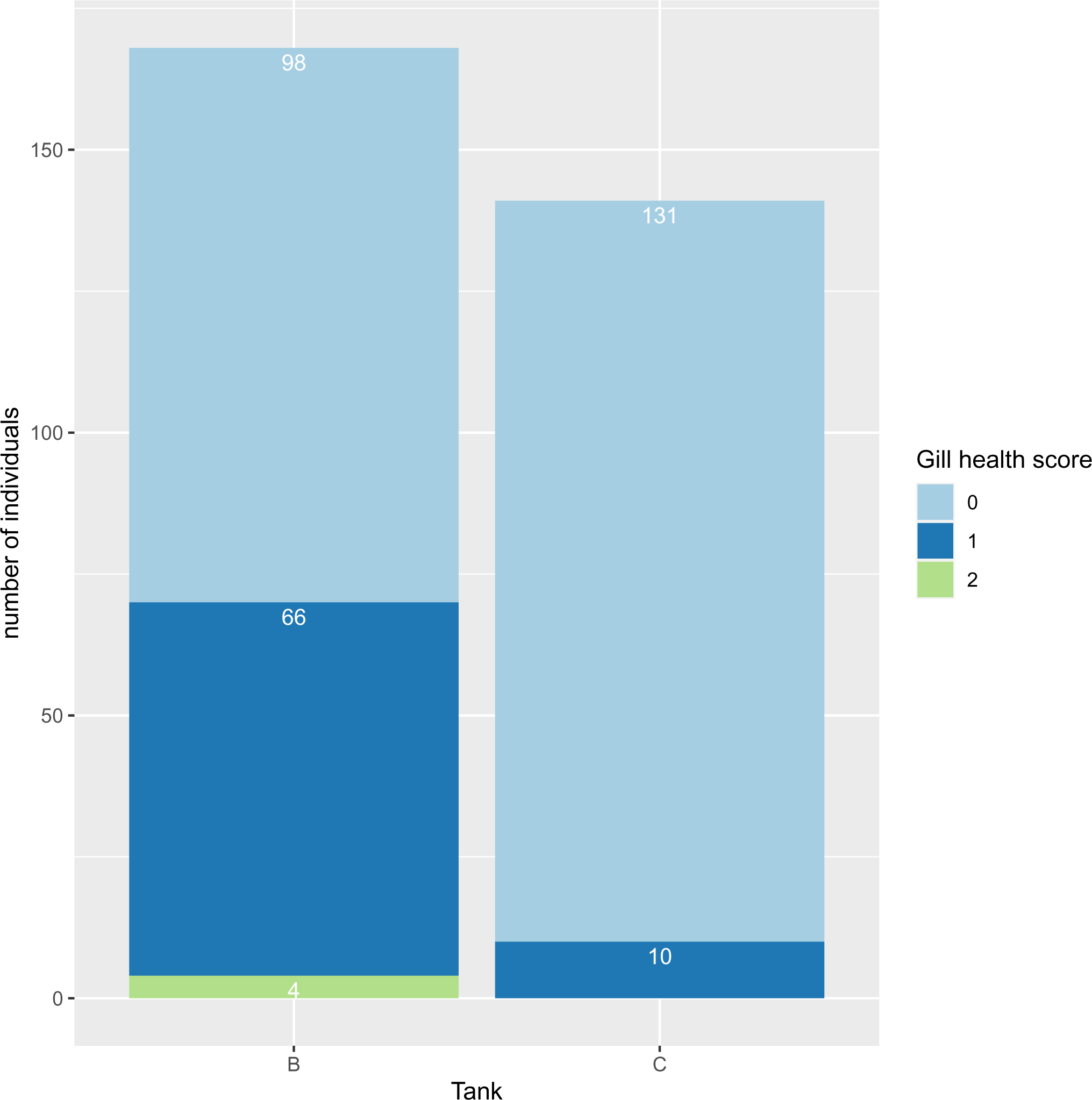
Bar plot of gill health scores by tank, where n the number of individuals

#### 3.1.4. QPCR assay

For most targets analysed, swabs were below Limit of Detection (LOD), with high Ct levels (Supplementary Information 2: Table 1). Ct values were in the quantifiable range for *Aeromonas hydrophila* and for the total bacterial load, with *Aeromonas hydrophila* being detected on all swabs (Supplementary Information 2: Table 2). *Aeromonas hydrophila* is a fresh water Gram negative bacterium, typically found in warmer climates. In addition, *Flavobacterium branchiophilum* was detected on some swabs, and although all signals were below the Limit of Quantification (LOQ), once presence of *Flavobacterium* is established, numbers are known to often rise rapidly.

### 3.2 Heritability estimates and genetic correlations

The genotyped fish originated from 54 full-sib, or maternal (3 dams contributed to more than one family) or paternal half-sib (10 sires contributed to more than one family) families, where family size ranged from 1 to 15 individuals, with a median of 10 individuals per family (Figure 11). The number of families in each tank is shown in Table 1; 51 families were represented across all tanks, and the number of offspring per family in each tank ranged from 1 to 6 (Tanks A and C), and from 1 to 5 (Tank B) offspring (Supplementary Information 3).

**Figure 11.**
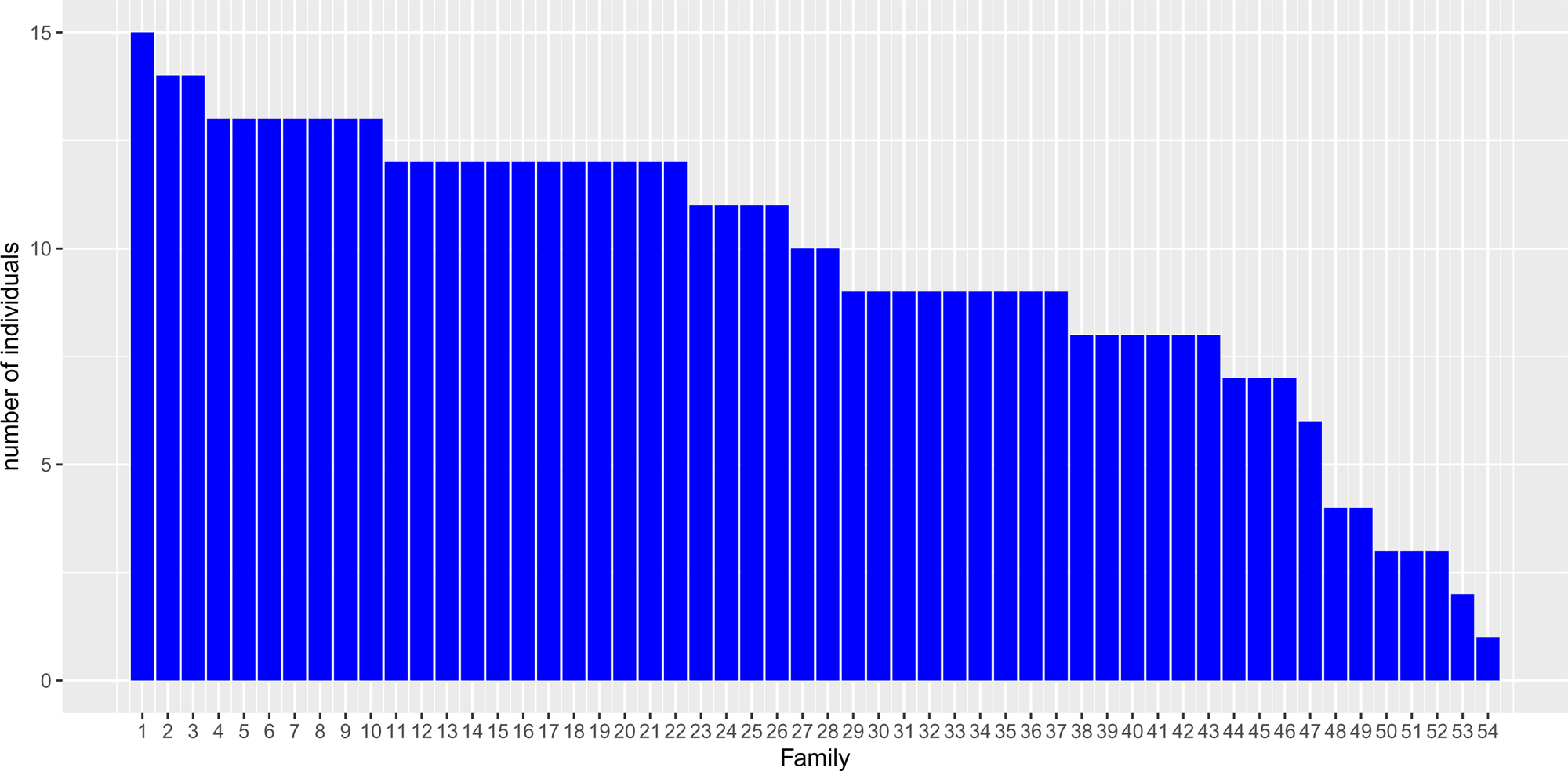
Histogram of family size in the data, where the Y-axis shows the number of individuals in each family

Genomic heritability estimates from the entire population were found to be ∼0.39 for body weight and length before and after challenge, and 0.26 for average daily weight gain (Table 4). The heritability for binary survival using a generalised linear model was found to be small and with large standard error. The continuous day-of-death trait using a linear model provided a moderately high heritability (Table 4), however, this result should be interpreted with caution due to the presence of truncated data with low prevalence of mortalities. Gill health score was analysed both as continuous and as binary (zero and non-zero score), however it was not possible to capture genetic variance for this trait in these data (77 fish had non-zero score and belonged to 42 different families), hence this trait could not be pursued further in the genetic analyses. Maturation (grilsing) was found to be heritable using a generalised linear model, which provided a heritability of 0.19. Using pedigree information rather than genomic relationships, provided rather inflated heritabilities for body weight, length and average daily weight gain, while for survival traits and maturation the estimates had large standard errors reflecting the relatively small number of families.

**Table 4.**
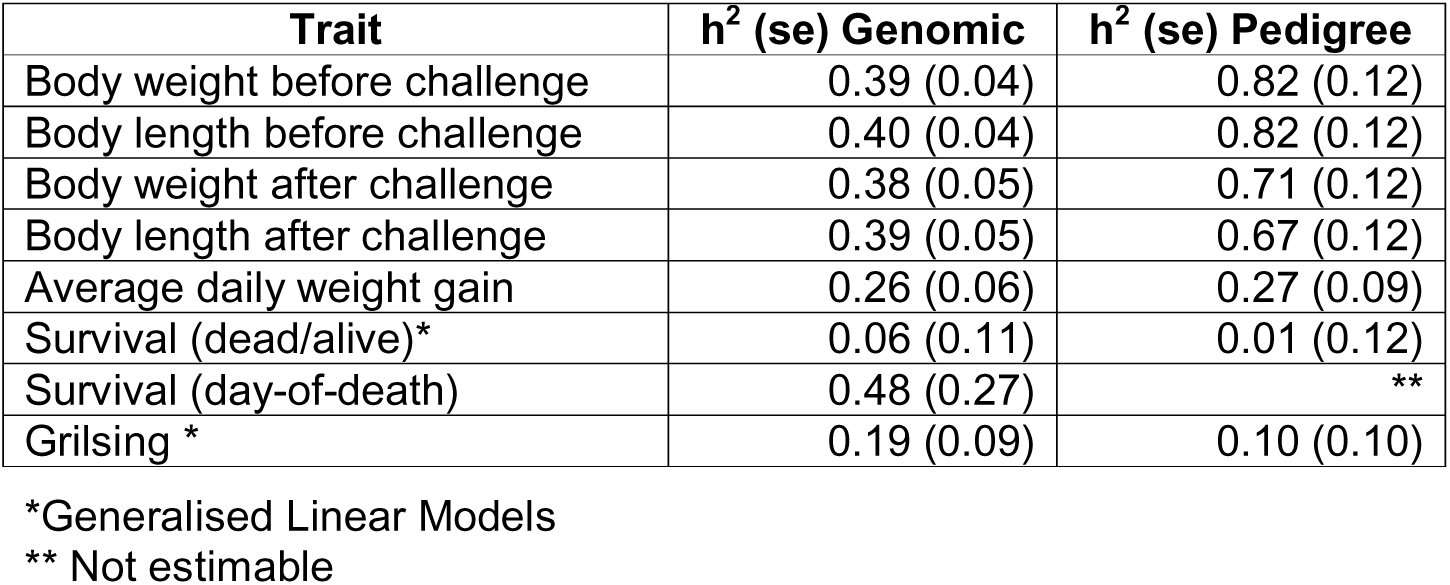
Heritability estimates obtained for each of the different traits and models, using genomic (i.e., genomic relationship matrix calculated from SNP marker data) and pedigree relationships.

Within-tank heritability estimates for body weight and length were within the same range as those observed for the overall population, being slightly lower after challenge (Table 5), and being lower for Tank C, which was likely due to the smaller number of individuals and families in this tank (Table 1) (for completeness, pedigree-based heritability estimates are shown in Supplementary Information 4). Genetic correlations between tanks for body weight and length before and after challenge were very close to 1, indicating no significant re-ranking between thermal environments in these data (Table 6). For average daily weight gain, heritability was substantially lower for the high temperature tank and very close to zero; this was reflected in the bivariate analyses so that genetic correlation could be estimated only between Tanks A and B, and was close to one, indicating no re-ranking of genotypes. For survival traits, heritability could be estimated only for the high-temperature tank as in Tank A all deaths occurred on day 29 and in Tank B the number of mortalities (n=5) was too small to provide sufficient information (standard error larger than the estimate for binary survival). Finally, for maturation, genetic correlations were very close to 1 but with large standard errors, and no re-ranking of genotypes was detected.

**Table 5.** Within-tank heritability estimates for each of the traits of interest using genomic relationships.

| Trait | $h^2$ (se) Tank A | $h^2$ (se) Tank B | $h^2$ (se) Tank C |
| --- | --- | --- | --- |
| Body weight before challenge | 0.51 (0.08) | 0.52 (0.10) | 0.36 (0.09) |
| Body length before challenge | 0.50 (0.09) | 0.53 (0.10) | 0.38 (0.09) |
| Body weight after challenge | 0.46 (0.09) | 0.39 (0.13) | 0.31 (0.11) |
| Body length after challenge | 0.48 (0.10) | 0.42 (0.12) | 0.40 (0.11) |
| Average daily weight gain | 0.34 (0.12) | 0.24 (0.14) | 0.03 (0.12) |
| Survival (dead/alive)* | ** | 0.11 (0.64) | 0.21 (0.15) |
| Survival (day-of-death) | ** | ** | 0.61 (0.25) |
| Grilsing * | 0.31 (0.24) | 0.15 (0.16) | 0.10 (0.19) |

\*Generalised Linear Models
\*\* Not estimable

**Table 6.** Genetic correlations between thermal environments using genomic relationships.

| Trait | Tanks A and B | Tanks A and C | Tanks B and C |
| --- | --- | --- | --- |
| Body weight before challenge | 0.98 (0.19) | 0.98 (0.19) | 0.99 (0.19) |
| Body length before challenge | 0.98 (0.18) | 0.99 (0.18) | 0.98 (0.20) |
| Body weight after challenge | 0.99 (0.22) | 0.98 (0.21) | 0.99 (0.28) |
| Body length after challenge | 0.97 (0.24) | 0.88 (0.18) | 0.88 (0.25) |
| Average daily weight gain | 0.99 (0.30) | ** | ** |
| Grilsing | 0.94 (0.85) | 0.95 (0.67) | 0.97 (1.60) |

\*\* Not estimable

## 4. Discussion

A moderate increase of average water temperature was found to be beneficial for growth compared to current ambient summer temperatures in Scotland, but even short-term exposure to more extreme heat-stress conditions had negative impacts. Fish in Tank B performed better than fish in Tank C, suggesting that increasing temperature beyond a certain threshold reduces performance. It was observed that even such short-term exposure to heat stress can be sufficient to generate significant effects on survival, and induce earlier maturation. Given that in the present study all fish in all thermal environments received the same amounts and type of feed, one possible explanation for the observed effects can be that those are due to the additional energy requirements of the fish exposed to higher temperatures in order to cope with more severe heat-stress. However, this study was performed within a commercial farm, hence having replicates for each of the thermal environments was not possible. This limits the statistical power to draw concrete conclusions, and the observations reported in this data need to be further tested in bigger experimental studies and larger populations. Nevertheless, the patterns observed here can inform the experimental design and choice of traits to monitor in future studies.

This study had a local character and a focus on effects from short-term temperature spikes. The aim was to observe effects under heat-wave conditions as they are expected to occur specifically at the west coast of Scotland. Therefore, it might not be possible to directly generalise these findings to observations from other geographical regions, where fish experience more prolonged heat-stress; coping mechanisms for prolonged heat-stress may be different at the biological and possibly also at the genetic level, compared to short-term responses to heat-waves. Previous studies have demonstrated that general trends obtained via global climate projections may over- or underestimate local effects, and in order to devise meaningful mitigation plans, finer-resolution predictions are needed (Falconer et al. 2020). Understanding heat-wave effects at the local level and in combination with existing on-farm practices, will allow us to better tailor farm and breeding programme management to more accurately reflect future local environmental conditions due to climate change.

The higher gill scores recorded, indicate poorer gill health due to higher water temperature, which is consistent with previous reports of compromised gill health and increased risk for disease outbreaks, for example amoebic gill disease (AGD) outbreaks when temperatures rise above 12°C (Benedicenti et al. 2019). We detected *Aeromonas hydrophila* on all swabs in the higher temperature tanks. *A. hydrophila* is a Gram negative aquatic bacterium of the genus *Aeromonas*, and is typically found in higher numbers in water temperatures > 10°C. This opportunistic pathogen causes a range of pathologies from skin infections to systemic disease known as motile aeromonad septicaemia, and can lead to outbreaks or sporadic cases when the right conditions converge. *A. hydrophila* can be responsible for substantial economic losses due to increased mortality and secondary infections. Vaccines against *A. hydrophila* are available, but mainly used in warm water aquaculture, where the sporadic outbreaks are more common. *A. hydrophila* is a zoonotic pathogen that can be responsible for potentially serious infections in humans through food poisoning or through aquaculture facilities (Abd-El-Malek 2017; Hayatgheib et al. 2020), therefore being a public health hazard, especially for people working in the aquaculture sector. In addition, recent studies report an alarming increase of antibiotic resistance of *A. hydrophila* against a number of commonly used antibiotics (Nhinh et al. 2021). The potential impact of a warming climate on the global distribution and prevalence of this pathogen requires further investigation. Emergence of temperature-dependent water-borne diseases in more northern latitudes due to global warming, has been reported for several pathogens; for example, emergence of *Vibrio cholera* infection in shellfish (El-Sayed and Kamel 2020), and increased risk for outbreaks of currently exotic diseases such as epizootic haematopoietic necrosis and epizootic ulcerative syndrome (Marcos-López et al. 2010). In addition, an increase of frequency and prevalence of endemic diseases may also be expected, such as furunculosis, proliferative kidney disease and white spot in salmonids, or koi herpesvirus in carp (Marcos-López et al. 2010).

Understanding the effect that heat-waves arising from climate change can have on salmon production can help us adapt the breeding goals locally, according to future environmental scenarios, aiming to breed fish that respond better to environmental stressors and climatic warming. The genetic analyses in this study aimed to explore to what extend phenotypic differences observed between different thermal environments, are controlled by genetics, and hence, to what extend it would be possible to use selective breeding to mitigate these effects. However, the number of records within individual tanks was small, with a substantial number of families having only 1 or 2 offspring within each tank (Supplementary Information 3), and estimates had large standard errors. Therefore, these results should be interpreted with caution, subject to further validation in larger populations. In addition, there was very low prevalence of grilse and of gill health scores > 0, while for some traits, there was not sufficient genetic variation in this population. This was reflected in the bivariate analyses performed to estimate covariances between thermal environments. For body weight, length, and average daily weight gain, there were no genotype-by-environment interactions detected in these data, and genetic correlations between tanks were very close to 1, indicating no significant re-ranking between thermal environments. However, a larger population would be needed to harvest more genetic variation within environments for a range of commercially important traits, and to be exposed to more severe and prolonged heat-stress conditions in order for the fish to fully express their genetic responses. Especially for disease-related traits, a disease challenge would be suitable in order to explore the combined impacts of disease and water temperature.

Site-specific temperature conditions need also to be considered in relation to changes in dissolved oxygen. Notably, higher temperatures not only increase the metabolic rates, hence requiring more oxygen, but also reduce oxygen solubility (Stehfest et al. 2017). As discussed in Lefevre et al. (2021), there are several hypotheses about the physiological mechanisms underlying fish responses to temperature change, including the gill oxygen limitation, and the oxygen- and capacity-limited thermal tolerance models; however, further studies are needed to provide the evidence base and the data to support those hypotheses and interpret the observed impacts. It is known that salmon express natural avoidance behaviour of both low dissolved oxygen and high temperature via natural movement within the water column (Stehfest et al. 2017). Therefore, strategies to optimize stocking density while sustaining profitability will be crucial in order to avoid overcrowding, and ensure fish welfare.

## 5. Conclusions

In conclusion, short-term exposure to a moderate increase of average water temperature may be beneficial for growth compared to current ambient summer temperatures in Scotland, but further exceeding salmon’s optimal zone will have negative impacts on growth, survival and can induce earlier maturation. In addition, poorer gill health shall be expected, accompanied by increased prevalence of pathogens currently found in warmer climates, such as *Aeromonas hydrophila*. Coping mechanisms for prolonged heat-stress may be genetically different to responses to short-term temperature spikes, hence, although these data did not reveal genotype-by-environment interactions, the potential for genetic mitigation needs to be further explored in larger populations exposed to more severe and prolonged heat-stress. It is likely that a combination of management solutions, genetic improvement, and optimized nutrition and feeding practices will be needed in order to plan effective mitigation strategies.

## Acknowledgements

The Authors gratefully acknowledge funding via the BBSRC Flexible Talent Mobility Award (BB/S50791X/1) and BBSRC Impact Accelerator Award (BB/S506722/1). ST would like to thank Professor Ross Houston (the Roslin Institute) and Sian Ringrose (Edinburgh Innovations) for their guidance and support to this project. The study was performed on The Scottish Salmon Company / Bakkafrost salmon strain. The authors would like to thank Robbert Blonk (Hendrix Genetics Aquaculture), and all the staff at the Ormsary Hendrix Genetics hatchery who performed the fish husbandry and data recording. ST would also like to thank Dr Carolina Peñaloza for her help with handling and storing gill swabs, and Dr Agustin Barria Gonzalez for the stimulating discussions on genetic analysis techniques and software.

## Author Contributions

Conceptualization and methodology: S. Tsairidou, A. Hamilton, J. van den Berg. Project administration and resources: J. van den Berg, S. Tapping, A. Hamilton. Formal analysis, software and funding acquisition: S. Tsairidou. Data curation, materials/analysis: S. Tapping, H. Sobolewska, A. Hamilton. Writing - original draft: S. Tsairidou; review & editing: A. Hamilton. All authors have agreed to the published version of the manuscript.

## Statements & Declarations

The authors have no competing interests to declare that are relevant to the content of this article.

## Data availability

The data in this study are not openly available due to their commercial nature.

## Supplementary Information captions

**Supplementary Information 1.** Structure analysis using classical multi-dimensional scaling: top left: all individuals in the study; top right: different colours indicate different families; bottom left: pink triangles represent female individuals and purple diamonds represent male individuals; bottom right: green triangles represent individuals in Tank A, black squares represent individuals in Tank B, and pink circles represent individuals in Tank C

**Supplementary Information 2.** QPCR results with Ct values and estimated target copy number for all pathogens tested (Table 1), and specifically for *Aeromonas hydrophila* and total bacterial load (Table 2)

**Supplementary Information 3.** Distribution of number of offspring per family within each of the tanks

**Supplementary Information 4.** Pedigree-based within-tank heritability estimates

