## Supplementary figures and images for "Assessing the effects of acute temperature changes of sea water temperature due to climate change on Scottish farmed Atlantic salmon and investigating genetic mitigation"

### Supplementary Information 1

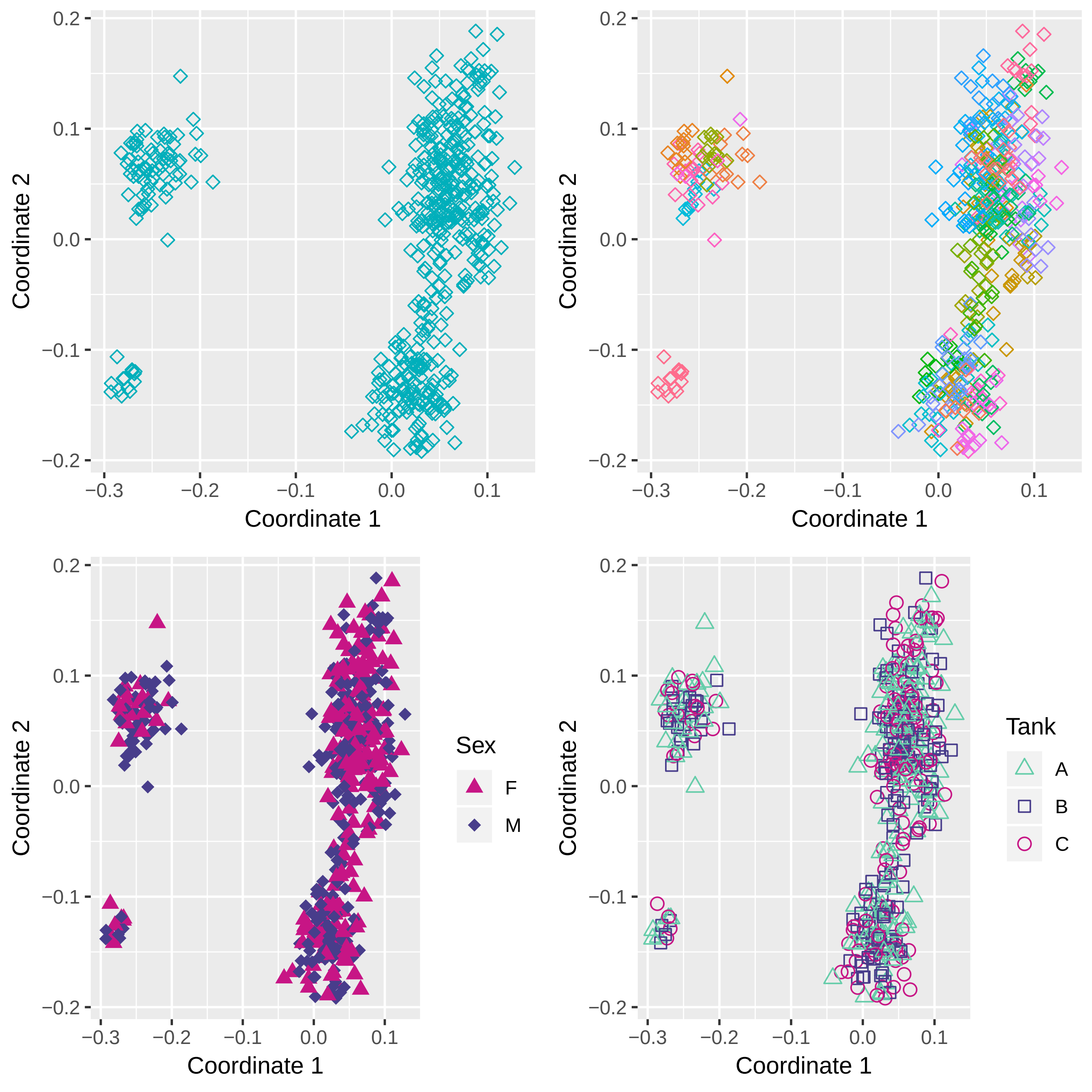

### Supplementary Information 3

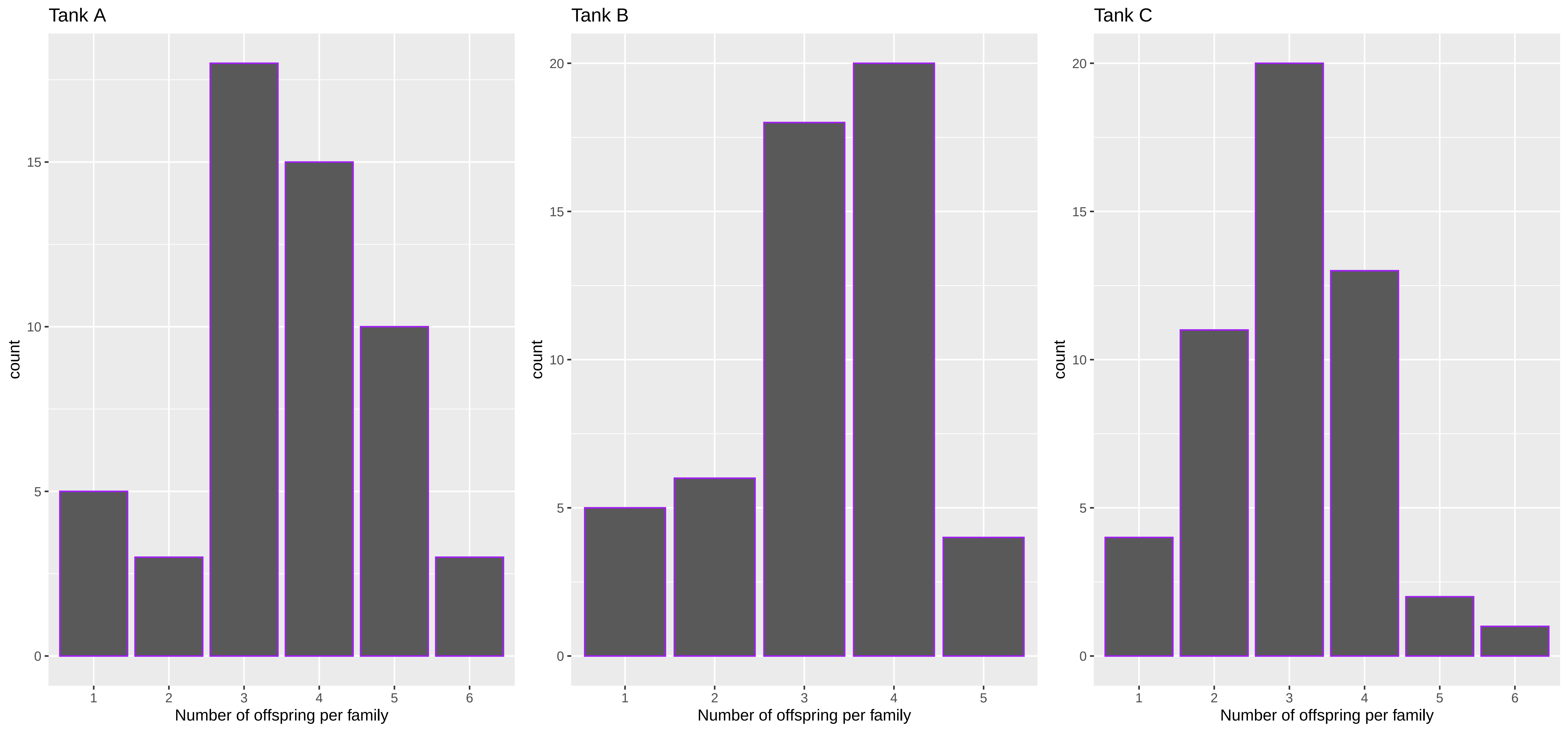
