## Supplementary Information 4 for "Assessing the effects of acute temperature changes of sea water temperature due to climate change on Scottish farmed Atlantic salmon and investigating genetic mitigation"

Within-tank heritability estimates and standard errors, for each of the traits of interest, calculated using pedigree relationships:

| **Trait** | **h^2^ (se) Tank A** | **h^2^ (se) Tank B** | **h^2^ (se) Tank C** |
| --- | --- | --- | --- |
| Body weight before challenge | 0.73 (0.17) | 0.59 (0.19) | 0.86 (0.18) |
| Body length before challenge | 0.77 (0.17) | 0.72 (0.19) | 0.83 (0.19) |
| Body weight after challenge | 0.66 (0.17) | 0.56 (0.20) | 0.61 (0.20) |
| Body length after challenge | 0.56 (0.17) | 0.61 (0.19) | 0.67 (0.20) |
| Average daily gain | 0.28 (0.17) | 0.46 (0.19) | * |
| Survival (dead/alive)* | * | 0.06 (0.72) | 0.11 (0.19) |
| Survival (day-of-death) | * | 0.09 (0.74) | * |
| Gill health score | NA | * | 0.05 (0.18) |
| Grilsing ** | 0.16 (0.33) | 0.06 (0.20) | 0.12 (0.19) |

* Not estimable

**Analysis performed using Generalised Linear Models
